# Trained Planaria Retain Memories Through Head Regeneration: A Model System for Insights into Non-Neural Memory and Neurodegenerative Diseases

**DOI:** 10.64898/2026.08.10.741213

**Authors:** Nikhil Dev, Angelina Nguyen, Michael Levin

## Abstract

Planaria exhibit remarkable regenerative ability, including the capacity to regrow complete heads and brains after decapitation. Here, we re-investigated whether regenerated planaria can preserve learned avoidance behavior, a phenomenon that has been reported previously but has been difficult to study due to unreliable experimental protocols. Using a light-to-food associative conditioning paradigm, planaria were trained to override their normal photophobic preference and then decapitated. Following a two-week regeneration period, behavioral responses to the conditioned stimulus were re-evaluated. Results indicated that the majority of regenerated planaria retained the learned response, supporting a model in which behavioral patterns can regenerate as well as anatomical patterns. By establishing a consistent, low-cost, and effective protocol for studying memory persistence through regeneration, such work may help inform future research on memory loss, resilience, and recovery in neurodegenerative diseases.

## INTRODUCTION

Planarians are freshwater flatworms renowned for their regenerative capacity. Small fragments of planaria can regrow into a full organism; in particular, planaria can generate an entirely new head and central nervous system after decapitation [1]. This is made possible by an abundant population of pluripotent stem cells (neoblasts) that continually replace tissues and allow body pattern restoration [2, 3]. Beyond regeneration, planarians have also been used as subjects for learning and memory research. Earlier experiments by James McConnell and others in the 1950s-60s suggested that planaria were capable of long-term memory [4–15]. For example, planarians could be conditioned to respond to light or vibrations and appeared to retain these memories even after regeneration. These studies raised fundamental questions about where and how memory is stored in these organisms. If a planarian can remember training after generating a new head, perhaps memory is not exclusively located in the synaptic circuits of the brain.

Memory storage in biological systems is commonly thought to occur via changes in synaptic connections among neurons [16], although other models are beginning to rise [16–22]. In different organisms, memory formation and recall are thought to be performed by the brain’s neural circuits (e.g., the hippocampus and cortex in mammals) and synaptic plasticity. In vertebrates, traumatic brain injury or neurodegenerative disease often results in permanent memory loss. For example, Alzheimer’s disease (AD), the most common form of dementia, exemplifies this problem: the disease contains extracellular amyloid plaques, intracellular tau tangles, and a profound loss of synapses and neurons, all correlating with a severe cognitive impairment and memory loss [23]. Current treatments for Alzheimer’s disease are very symptomatic and fail to address the underlying neurodegeneration. The human brain has very limited regenerative ability - only very specific stem cells seem to be able to produce new neurons (in the hippocampus and olfactory bulb), and this neurogenesis declines with age and in Alzheimer’s patients. Similarly, damage to the human brain usually results in permanent memory loss, since neurons and the information they hold can often not be recovered.

Animals which regenerate brain tissue provide an important opportunity to test assumptions about the location and mechanism of memory, and the potential transfer of memories across tissue in vivo [24] which could have massive implications for design of, and consequences of, regenerative therapeutics in human brains targeting aging, disorders of memory, and degenerative disease. Planaria are a model system in which learning and brain regeneration can be done in the same animal, but their behavioral individuality has driven the continued search for robust, low-cost protocols for memory regeneration, accessible to all stages of researchers. Here, we re-examine the capacity of planarian flatworms to retain a specific learned behavior (a trained light preference) after decapitation and regeneration of the head.

### Classical data: a brief literature review

Early reports of planarian learning date back over half a century (reviewed in [6, 7, 24]). In those studies, planarians were taught tasks such as navigating mazes or associating light with shock, and surprisingly, some worms appeared to retain these memories after being cut and regenerated. More rigorous modern experiments have confirmed that planaria can form long-term memories and retain them through regeneration. Using automated methods, Dugesia japonica planarians were trained in an automated apparatus to prefer a rough-textured, illuminated surface (a situation they would normally avoid) by pairing the lit area with food reward [25]. After two weeks of training, the worms overcame their light avoidance. The researchers then amputated the planaria’s heads, removing the brain and sensory organs. Approximately 14 days later (the time required for full regrowth of the head and brain), the planarians were tested for memory of the training. The decapitated worms that had regenerated their heads demonstrated significantly faster re-learning (savings) and greater liking for the light-associated context compared to control worms that had never been trained. In other words, the regenerated worms *“exhibited evidence of memory retrieval in a savings paradigm after regenerating a new head”.* This provided clear, quantitative evidence that some memory of the prior training survived the complete destruction and replacement of the original brain [7]. The authors proposed that planaria represent a powerful model to investigate *“the fundamental interface between body patterning and stored memories,”* since these animals seem to encode memories in a way that can connect with a newly built nervous system. Our present experiment builds on this work, using a similar conditioning paradigm to probe memory retention in regenerated planaria.

How might planaria store memories outside of the brain? One possibility is that residual neural tissue in the body (for example, peripheral neurons or the nervous system in the tail stump) holds a memory trace that can educate the new brain. Another possibility is that memory is encoded in non-neural cells by lasting molecular changes. Epigenetic memory has gained popularity as a concept, especially given evidence from other invertebrates. Insects provide a strong example: in a study by Blackiston et al. (2008) [26], caterpillars of the moth *Manduca sexta* were trained to avoid a particular odor by pairing the odor with a mild shock. After metamorphosis into adult moths (a process that disintegrates and reorganizes the caterpillar’s body and brain), the moths were tested and found to still avoid the odor they had learned to dislike as larvae. This was a demonstration that associative memory can survive metamorphosis in Lepidoptera, implying that despite the remodeling of the nervous system, some cellular or molecular record of the learned behavior persisted and influenced the rebuilt brain [26] (see also [27–30]). The authors speculated that the memory might be preserved in a subset of neurons that are carried over or by conserved molecular signals (such as modifications to neurons’ DNA or proteins) that guide the reconnection of the circuit in the adult [7].

Recent research on memory transfer via RNA provides another angle on non-synaptic memory storage. Bédécarrats et al. induced a simple form of long-term memory (sensitization of the defensive withdrawal reflex) in Aplysia by applying tail shocks [31]. Then they extracted RNA from the nervous systems of these “trained” animals and injected it into untrained snails. Amazingly, the recipient animals began to exhibit reflex responses as if they had undergone the same training, whereas control Aplysia injected with RNA from untrained donors showed no such change [31]. The transfer effect appeared to depend on epigenetic mechanisms: the injected RNA triggered increased DNA methylation in the recipients’ neurons, and blocking DNA methylation prevented the behavioral change [31]. These findings suggest that RNA from trained animals carried specific information which could “induce an epigenetic engram” in recipient animals. This work possibly revives the idea that some components of memory (a priming of neural excitability or gene expression states) can be stored in molecules like RNA and are not locked within the synapse. Such changes might persist in planaria after decapitation, e.g., in the remaining tissue and instruct the newly developed neurons, essentially imprinting the new brain with the memory.

### Current Context: Neurodegenerative Disease Research

In human systems, the idea that memory could be preserved through neuronal turnover or stored in chemical form is not a popular part of neuroscience, mainly because humans do not naturally replace entire brain regions. Nevertheless, research in cognitive disorders is exploring more treatments beyond synaptic pharmacology, including regenerative medicine and epigenetic therapy. One major area of interest is adult neurogenesis. The adult human hippocampus generates new neurons, although the rate decreases with age. Studies have observed impairments in this neurogenic process: the neural stem cell niches produce fewer new neurons, and those that are born often struggle to survive or integrate into the environment. Epigenetic alterations, such as hypermethylation of pro-neurogenesis genes and excessive histone deacetylation in human brains, can suppress factors essential for neural stem cell proliferation and differentiation. Reversing these epigenetic blocks has shown good results in experimental models [32]. Studies reviewed that treatment with histone deacetylase inhibitors (like valproic acid) or other epigenetic modulators can “promote neuronal generation” in the adult brain and improve cognitive outcomes in models, essentially by reactivating developmental programs for neuron growth [33]. Another strategy is direct stem cell transplantation. Preclinical trials in transgenic neurodegenerative mice using mesenchymal stem cells or induced neural progenitors have reported several benefits: cell factors that reduce neuroinflammation, support the health of existing neurons, and even differentiate to replace some lost cells. These actions have improved spatial memory performance in treated AD-model mice relative to untreated controls [34]. Early-phase clinical trials (for example, using umbilical cord blood-derived stem cells injected into patients’ brains) have shown safety and potential cognitive improvements, though efficacy remains to be proven. Importantly, any regenerative approach in humans faces the challenge that new neurons would need to form meaningful connections to recover memories or functions lost – something we do not know how to achieve yet. This is where understanding natural models of brain regeneration, like planaria, becomes highly relevant. If we can determine the signals or molecules by which a planarian’s body instructs a new brain to take on old memories, similar signals might be produced or mimicked to help integrate new neurons in the adult brain. For instance, planarian studies might reveal memory-specific mRNA or epigenetic markers that ensure continuity of memory through regeneration. Those markers (be they transcription factors, noncoding RNAs, or epigenetic modifications) could enhance synaptic re-connection or circuit reassembly in the injured or diseased brain.

In summary, prior research shows that several model systems offer evidence of memory persisting across significant regenerative and repair events. Alzheimer’s disease research could be strongly potentiated in new directions by reaching beyond neuron-centric approaches. By confirming and memory retention in regenerating planaria, accessible planarian protocols contribute to a roadmap seeking an understanding of memory that transcends mature neurons and toward a broader reparative function for patterns of form, behavior, and physiology. Such understanding could eventually form strategies to protect or restore memories in human neurodegenerative diseases and regenerative medicine more broadly.

## METHODS

### Planarian Subjects

Our study used freshwater planarians obtained from a laboratory colony (mixed species of *Dugesia* or *Girardia*, ∼1–1.5 cm length). All worms were maintained in clean spring water at room temperature (∼20°C) and fed organic beef liver twice weekly. Before training, worms were starved for 5 days to ensure high motivation for food during the conditioning trials.

### Experimental Design

The experiment consisted of three phases: (1) baseline preference assessment, (2) training/conditioning, and (3) post-regeneration memory test – conducted on the same group of planaria. A total of 34 worms were used for group preference tests, and a subset of 10 individuals from this group were tracked longitudinally for detailed analysis of their behavior before and after regeneration.

### Apparatus and Behavioral Measures

To assess planarian light/dark preference and learning, we used a Petri dish (9 cm diameter) divided into equal light and dark regions. One half of the dish was illuminated from above by a UV Light (365+395nm wavelength, 5V 3W), and the other half was kept dark by an opaque cover. During trials, each worm was placed in the center of the dish, and its position (light or dark side) and movement were recorded. For group preference assays, multiple worms (up to 34) were tested simultaneously in a larger divided tray, and counts of how many worms were on the light side vs. dark side were taken at 2-minute intervals. For individually tracked worms, we recorded the time each worm spent on the light side over a 10-minute trial (using video tracking and manual stopwatch verification).

### Baseline Preference Test

Prior to any training, we measured the innate preference of planaria for light vs. dark. It is well-documented that planaria exhibit photophobic behavior. They naturally prefer dark environments. In our baseline trial, 34 worms were placed in the dish and allowed to move freely for 30 minutes, with positional counts taken every 2 minutes. As expected, the planaria strongly favored the dark half; at the 2-minute mark only 6 of 34 worms (∼18%) were on the illuminated side, and throughout the trial the majority remained in darkness (e.g., at 10 minutes, 9 on light vs. 25 on dark; at 30 minutes, just 3 on light vs. 31 on dark) (see Supplement, Table A2). This showed a strong natural dislike of the light side.

### Training Protocol

We conditioned the planaria to associate the illuminated side of the arena with a food reward, thereby attempting to override their natural aversion. Training took place over 10 days with one session per day. In each session, worms were placed in the divided dish with the light on one side. A 1 oz intact piece of fresh beef liver (a highly attractive food for planaria) was placed on the illuminated side. Worms were allowed to locate and consume the food over a 15-minute period. To encourage exploration, tapping was applied if a worm stayed in the dark quadrant for more than 5 minutes; this was rarely needed as hungry worms eventually wandered in search of food. By repeatedly experiencing food in the light zone, the planaria were expected to form an association that mitigated their photophobia. The exact positions of the worms during training were not quantitatively recorded, as the presence of food alters natural exploratory behavior and causes worms to aggregate around the food source. However, by the end of training, we observed that worms easily moved into the lit area when placed in the dish, showing they had learned to change their behavior.

Twenty-four hours after the final training session, we conducted a post-training preference test identical to the baseline assessment. The same 34 trained worms were placed in the dish (now with no food present), and their distribution was recorded every 2 minutes for 30 minutes. The results showed a dramatic shift from the baseline: after training, the majority of planaria now preferred the light side. At 2 minutes into the test, 30 of 34 worms (∼88%) were found on the illuminated side, with only 4 remaining in darkness. This strong preference persisted; for instance, at 10 minutes, 19 were on light vs. 15 on dark, and by 30 minutes, 33 of 34 worms were on the light side (Supplement Table A1). Thus, the conditioning was successful in reversing the innate behavior – the worms learned that the lit side was associated with a positive outcome (food) and therefore spent much more time in light than other worms would. This demonstrated long-term memory of the training, as the test took place a day after the last training trial, with no food present.

### Surgery and Regeneration

Immediately following the post-training test, we proceeded to amputate the heads of the trained planaria to begin the regeneration phase. Using an exacto knife on a cooled glass plate, each worm was decapitated by making a cut just posterior to the pharynx (approximately at the planarian mid-body). This method cleanly removes the head, including the brain (located in the anterior dorsal region), while leaving a sizable tail piece. The head portions were discarded. The tail fragments (which, for clarity, we term “decapitated worms”) were transferred to fresh spring water and maintained in individual dishes. No feeding was done during regeneration to avoid new learning; planaria can survive weeks without food and instead rely on their internal stores during regrowth. The water was changed every other day. Over the next 14 days, the tail fragments regenerated their missing anterior structures. By day 7 post-amputation, eye spots were visible in all regenerating worms, and by day 14, each worm had regrown a complete head that looked just like the original. We thus scheduled the memory retention test at two weeks post-decapitation, allowing ample time for neural circuits to reform. It is important to note that during the regeneration period, worms were kept in a neutral, low-light environment with no explicit exposure to the experimental light/dark dish or any food rewards – this was to ensure that any behavior observed in the upcoming test could be attributed to retained memory rather than new learning during regeneration.

### Post-Regeneration Memory Test

To evaluate memory retention, we subjected the regenerated planaria to the same conditions as the earlier preference tests. Each worm (now with a brand-new head/brain) was placed in the light/dark dish with no food present, and behavior was measured over 10-minute trials. Because of the smaller sample (some worms did not survive the full two weeks; we ended up with 30 viable regenerated individuals out of the original 34), and to allow pairwise comparison, we focused on tracking a subset of 10 worms individually through repeated trials. We conducted three trials for each worm on consecutive days (days 14, 15, and 16 post-amputation) to assess consistency. During each 10-minute trial, we recorded the total time (in seconds) the worm spent on the illuminated side. This metric served as an indicator of its preference or aversion to light – longer time in light suggests the worm finds the light side attractive (as in a trained worm), whereas short time in light suggests it is avoiding light (as an untrained worm would). For additional comparison, we also performed parallel testing on a control group of worms that had been decapitated and regenerated without prior training. However, we emphasized the within-subject comparison of each trained worm before and after regeneration.

## Data Analysis

The key comparisons of interest were: (a) group-level light-side preference of trained worms before vs. after regeneration, and (b) within-subject light-side time for each worm with its original brain vs. with its regenerated brain. For the group preference counts, we looked at the proportions of worms on the light side at each time point and across the trial duration. For the individual time-in-light data, we calculated the mean time in light (out of 600 seconds) for each worm pre- and post-regeneration (averaging its three trials in each condition). We then computed the overall mean ± standard deviation of these times for the original-brain trials and the regenerated-brain trials, and used a paired statistical test (two-tailed paired t test) to check for any significant difference, with significance threshold set at *p* < 0.05. We also examined whether any “relearning” occurred during the three post-regeneration trials by looking for trends (e.g., did time in light increase from trial 1 to trial 3 post-regeneration, which might indicate the worm had partially forgotten and was re-acquiring the preference).

## RESULTS

### Training Induced a Reversal of Innate Light Aversion

Before conditioning, planarian worms strongly avoided the illuminated side of the environment. Over a 30-minute baseline trial, the vast majority of worms stayed in the dark region (e.g., at 2 minutes, only 6/34 worms were in light; at 30 minutes, 3/34 were in light), confirming an inherent photophobic behavior (Supplement Table A1). This baseline distribution was significantly different from chance expectation (50% light/50% dark), with light occupancy significantly lower than expected at both 2 minutes (binomial test, *p<0.01*) and 30 minutes (binomial test, *p<0.01*). After 10 days of training with food reward in the light zone, this preference was dramatically changed. In a test conducted one day post-training, the worms exhibited a pronounced preference for the light side (Supplement). At the 2-minute mark of the post-training trial, 30 out of 34 worms (∼88%) were on the illuminated side seeking the expected food reward, compared to only 4 remaining on the dark side. This post-training light occupancy was significantly greater than chance (binomial test, *p<0.01*). Additionally, the increase in light occupancy from baseline to post-training at 2 minutes (6/34 vs. 30/34) was statistically significant (chi-square test, *p<0.01*), supporting a reversal of the original avoidance response. This ratio remained skewed toward light throughout the trial (33/34 in light at 30 min; see Supplement). Light occupancy at 30 minutes was significantly higher than chance (binomial test, *p<0.01*), and the shift from baseline to post-training at 30 minutes (3/34 vs. 33/34) was also statistically significant (chi-square test, *p<0.01*). Thus, the conditioning protocol was successful in creating a long-term associative memory. This conclusion is supported by the significant baseline vs. post-training differences in light preference across the trial (chi-square tests, *p<0.01*). No such change was observed in untrained control worms (which continued to avoid light). Control worms remained significantly photophobic during testing, with light occupancy significantly below chance (binomial test, *p<0.01*), indicating that the preference reversal was not due to spontaneous behavioral drift. Trained worms also differed significantly from untrained controls in post-training light occupancy (chi-square test, *p<0.01*), further supporting that the observed effect was driven by conditioning rather than environmental factors. Finally, the trained worms’ behavior demonstrates long-term memory retention of the training, as they were tested after a 24-hour rest with no food present and still showed the learned preference. The persistence of significantly elevated light occupancy after the rest period (binomial tests, *p<0.01*) supports that the learned association was retained beyond the immediate training context and reflects long-term memory rather than short-term arousal or immediate reinforcement.

### Planaria Retain Learned Preference After Head Regeneration

The central question was whether the trained behavior would persist after the animals had their heads (and original brains) removed and regenerated. Following decapitation and a two-week regeneration period, we tested the same worms (now with regenerated brains) in the light/dark arena with no food cues. The regenerated planaria still showed a strong inclination toward the illuminated side, spending 7.65 ± 1.63 minutes in the light compared with 6.87 ± 1.63 minutes before regeneration (paired t-test, p > 0.05). Whereas never-trained planaria with regenerated heads would be expected to avoid light, our trained and regenerated worms spent substantial time in the light area, indicating retention of the training memory.

Additionally, we tracked 10 individual planaria through three trials before and after regeneration. With their original brains (trained), the 10 trained worms spent on average 6.87 ± 1.63 minutes (mean ± SD, n=15 trials pooled) out of 10 minutes on the light side, reflecting their learned preference for light. After head regeneration, the same worms averaged 7.65 ± 1.63 minutes (n=15 trials) on the light side. This corresponds to a slight increase in time-in-light post-regeneration. In other words, there was no detectable loss of the learned memory due to regeneration – if anything, some worms appeared to favor the light even a bit more with their new brains (the group mean increased by ∼47 seconds). Table 2A (Supplement) plots the before-vs-after times for each worm; most points lie near the diagonal (unity line), signifying consistency of behavior pre and post-regrowth. All regenerated worms moved toward the light side and spent an average of 7.65 minutes of the 10-minute trial in the illuminated region. Not a single regenerated worm showed the typical strong dark preference of worms, which previously spent less than 30% of time in light during baseline trials.

**Table 2.** Individual Planarian Time Spent in Light (seconds out of 600 s) in 10-Minute Trials Pre- and Post-Regeneration. Data for 10 representative worms are shown, each tested in three trials with their original brain (pre-regeneration) and three trials with their regenerated brain (post-regeneration). “Orig” = original brain trials, “Regen” = regenerated brain trials.

| Worm ID | Original brain<br>(Trial 1) | Regenerate<br>d brain<br>(Trial 1) | Original brain<br>(Trial 2) | Regenerate<br>d brain<br>(Trial 2) | Original brain<br>(Trial 3) | Regenerate<br>d brain<br>(Trial 3) |
| --- | --- | --- | --- | --- | --- | --- |
| 1 | 5 | 6 | 4 | 4 | 10 | 9 |
| 2 | 7 | 7 | 6 | 7 | 6 | 6 |
| 3 | 3 | 5 | 8 | 8 | 4 | 5 |
| 4 | 6 | 6 | 9 | 10 | 6 | 6 |
| 5 | 9 | 10 | 3 | 5 | 6 | 7 |
| 6 | 4 | 4 | 6 | 5 | 8 | 9 |
| 7 | 6 | 7 | 7 | 7 | 9 | 10 |
| 8 | 8 | 9 | 7 | 6 | 7 | 7 |
| 9 | 7 | 9 | 6 | 8 | 10 | 10 |
| 10 | 3 | 3 | 9 | 10 | 8 | 8 |

### Behavioral Observations

Apart from the time measurements, we noted qualitative behavior that supports memory retention. Regenerated planaria often turned toward and moved into the light area when placed in the test dish, showing behavior similar to what they learned instead of just exploring randomly. In contrast, untrained-regenerated control worms (which had never been trained but went through decapitation and regrowth) generally settled in the darker area, showing clear photophobia. The trained/regenerated worms also showed anticipation at the light/dark boundary, as if expecting food once they reached the lit side – some began probing the area where food used to be given during training, even though no food was present. This indicates that the associative memory (“light means food”) remained linked.

No significant differences in speed or general health were observed between regenerated worms and their pre-decapitation selves; regeneration did not impair their basic sensory or motor functions in any obvious way. All regenerated worms responded normally to stimuli and had intact negative phototaxis when tested in an untrained context (for instance, if removed to a novel environment, they would avoid bright light, except in the specific context of the training arena where they had learned otherwise). This reinforces that the effect we observed was specific to the learned association, not a generalized change in light sensitivity.

In summary, the results clearly show that planaria kept their learned light-seeking behavior even after their heads were cut off and regrown. The time they spent in the light was almost the same before and after regeneration, and much higher than that of untrained worms. This strongly supports the idea that planaria can retain memory through brain regeneration.

## DISCUSSION

We confirmed that planarian flatworms can retain a specific trained memory through the process of regenerating their brain. The study should be understood as a replication using a simplified and accessible conditioning protocol rather than an exact methodological replication of prior results. Our experiment used a manual light-food associative conditioning paradigm, relying on repeated exposure to food placed exclusively in an illuminated environment to change the planarians’ avoidance of light. This approach differs from earlier protocols by emphasizing direct positive reinforcement over extended daily training, rather than aversive stimuli or automated operant conditioning.

In our experiments, worms that learned to associate light with food continued to show the same preference after their heads were removed and regrown. The simplest interpretation is that whatever changes in the worm’s system encoded the memory of “light is good” were not entirely lost when the brain was removed. Instead, those changes must have resided in the remaining body or in some transferable form, and they influenced the assembly or function of the new brain such that the learned behavior was expressed nearly immediately upon regeneration.

It is still unknown where the memory resides, whether it involves a neural substrate, or whether existing cells actively “train” the new brain after it regenerates or whether the memory moves across tissue in a mechanism different from that used in normal learning. Given the symmetry between patterns of form and those of behavior, and conserved molecular mechanisms underlying both [35, 36], additional potential mechanisms could involve molecular and epigenetic memory. Planarian regeneration is made by positional cues and stem cell differentiation. If the cells in the stump carry biochemical markers of the experiences the worm had (such as phosphorylation of certain proteins, up- or downregulation of specific genes, or storage of particular RNAs), these could influence the properties of the new neurons as they form. For example, during training, the planarian’s body cells, such as muscle or skin cells, which produce many important developmental genes, might change how they express certain neurotrophic factors or guidance molecules in response to the new behavior. When the head regrows, these factors could influence how the new brain’s connections form or how excitable its neurons are, favoring the trained behavior. This idea is similar to recent RNA experiments in snails (Aplysia) [31, 37, 38], where behavioral changes linked to gene expression in neurons (like DNA methylation) were shown to transfer memory-like effects when RNA was injected into other neurons. This suggests that some long-term memories could involve an epigenetic “footprint” – a set of RNA and protein changes that help maintain altered neuron responses over time.

In planaria, there is evidence for such epigenetic involvement. Some modern studies have explored molecular explanations for the historical “memory transfer” observations. For example, experiments examining RNA interference and molecular signaling pathways suggest that RNA-based mechanisms could influence behavioral memory processes in planaria, though the phenomenon remains controversial [6]. Furthermore, planarian neurons themselves may undergo activity-dependent gene expression changes when learning, which then persist in the regenerating fragments. Our experiment cannot pinpoint the exact mechanism, but the fact that regenerated worms did not require extensive re-training suggests that the memory trace was not vague – it was precise enough to immediately bias behavior, hinting at a structured preservation of information.

### Implications for future medicine

What does this mean for more complex brains and specifically for neurodegenerative disease? While one must be cautious in extrapolating from flatworms to humans, memory could use highly-conserved mechanisms across evolutionary taxa and thus might be more resilient and distributable than traditionally thought. The prevalent view is that memory fails because synapses and neurons degenerate. Efforts to treat diseases thus far have focused on preventing neuron death or clearing pathological proteins, with limited success. Our findings are consistent with a different perspective: if even a simple brain can have its entire neural substrate replaced yet still recall information, perhaps memory is encoded in part by biochemical states that could, in principle, be revived or transferred. It invites speculation that even in an Alzheimer’s brain, some latent traces of lost memories might persist [39] – for example, in altered gene expression patterns of surviving neurons or glial cells [40] or perhaps cytoskeletal structures [41, 42]. If we could find ways to reactivate or reinforce those traces, maybe through epigenetic drugs or neurotrophic factor therapy, we might recapitulate portions of memory function. In fact, some studies have shown that HDAC2 overexpression in AD models leads to memory deficits because it overly silences genes needed for synaptic plasticity; conversely, blocking HDAC2 allowed those genes to be re-expressed and restored memory performance. This is analogous to “unlocking” a memory that was stored but not accessible. In planaria, one could argue the memory was stored in an inaccessible form (while the brain was absent) and became accessible again once the new brain formed – hinting that the information never truly disappeared but was lying dormant in the organism’s cells; perhaps the brain’s neural circuits have an interpretive function over a molecular substrate [43], in addition or instead of whatever storage mechanism they may have.

Another application is in regenerative medicine for the brain. Currently, replacing lost neurons (through stem cell grafts or by stimulating endogenous stem cells) is a hot area of research for neurodegenerative diseases, stroke, and spinal cord injury. A key challenge is that new neurons need to integrate into existing networks for meaningful recovery. Planaria provide a successful example of entire networks being regenerated and integrated such that old functionality (here, learned behavior) is restored. What can we learn from them? Perhaps the importance of the environment: planarian tissues likely provide instructive signals to guide new neurons. In mammals, the degenerated brain environment is hostile – full of inflammation, misfolded proteins, and lacking support signals – which may explain why new neuron survival and integration is poor. This suggests combining regenerative attempts with the modulation of the milieu. In an interesting mouse study, adding newborn neurons alone did not improve cognition in AD models, but adding them along with factors that enhanced the synaptic environment (BDNF, for instance) did yield cognitive benefits. Similarly, physical exercise in aged mice (which increases neurogenesis and releases systemic factors) improved memory, and even when those factors were transferred via plasma to other mice, they stimulated neurogenesis and cognition. These results echo the idea that a holistic approach – not just a cell transplant, but also tweaking the bioelectric, epigenetic, and trophic environment – could be needed for memory restoration. Planaria naturally do this: when a head regenerates, the body has already upregulated a host of developmental genes and established gradients to ensure proper wiring.

### Limitations of the study

Planarian memories that have been shown experimentally are much simpler than the full range of human memories – we trained a basic environmental association, not a complex episodic memory. That, in addition to the difference between planaria and mammals, suggests caution in extending the current data to clinical settings. However, we believe it likely that induction of regenerative repair in the brain would likely involve not only structural and functional, but also historical, patterns.

Another limitation is that we didn’t identify the memory mechanism in planaria. Future studies could try transplanting different sizes and pieces of tissue, or even single-cell transplant-based reconstitution [44], from trained worms to untrained ones, to see if memory transfers, or using drugs during regeneration to block specific gene activity or neural signals, as loss-of-function experiments to identify mechanism.

## CONCLUSION

Demonstrations of memory persistence in regenerating animals broaden our thinking about memory storage and recovery. While human brains are highly complex, exploring these alternative memory mechanisms could inspire novel approaches to treating memory loss. In Alzheimer’s disease, treatments that promote the growth of new neurons, strengthen synapses, or reactivate inactive gene programs could help restore some cognitive function, as could entirely orthogonal treatments that facilitate the movement of stored information across tissues. By combining these approaches, scientists and clinicians may one day develop therapies that not only halt neurodegeneration but also recover lost memories – achieving a longstanding goal inspired by a small flatworm.

**Figure 1:**
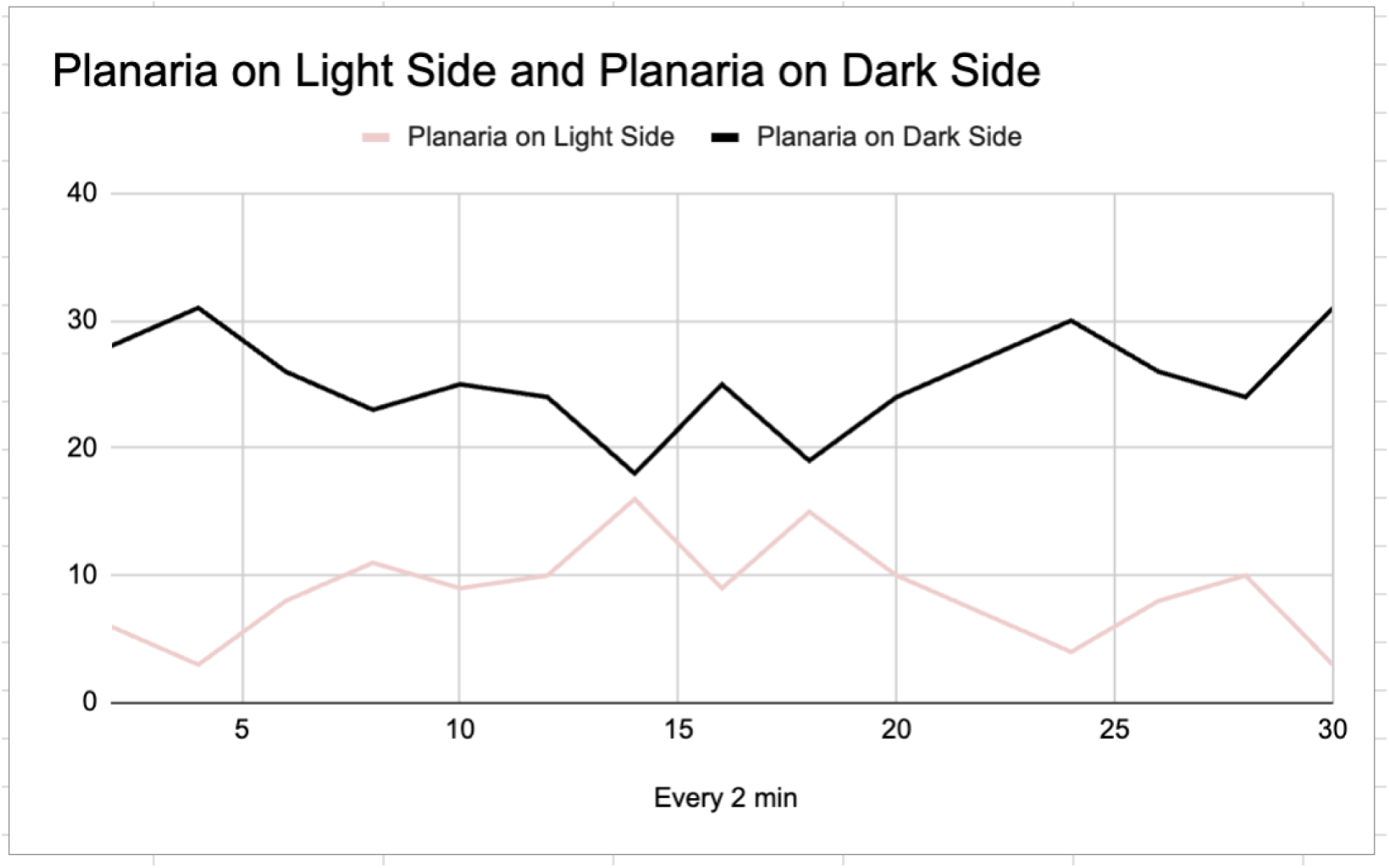
Pre-training preference of planaria between the light and dark sides of the test arena. Data show the number of planaria observed on each side every 2 minutes over a 30-minute period (n = 34). Prior to training, planaria predominantly preferred the dark side, indicating a natural baseline aversion to light.

**Figure 2:**
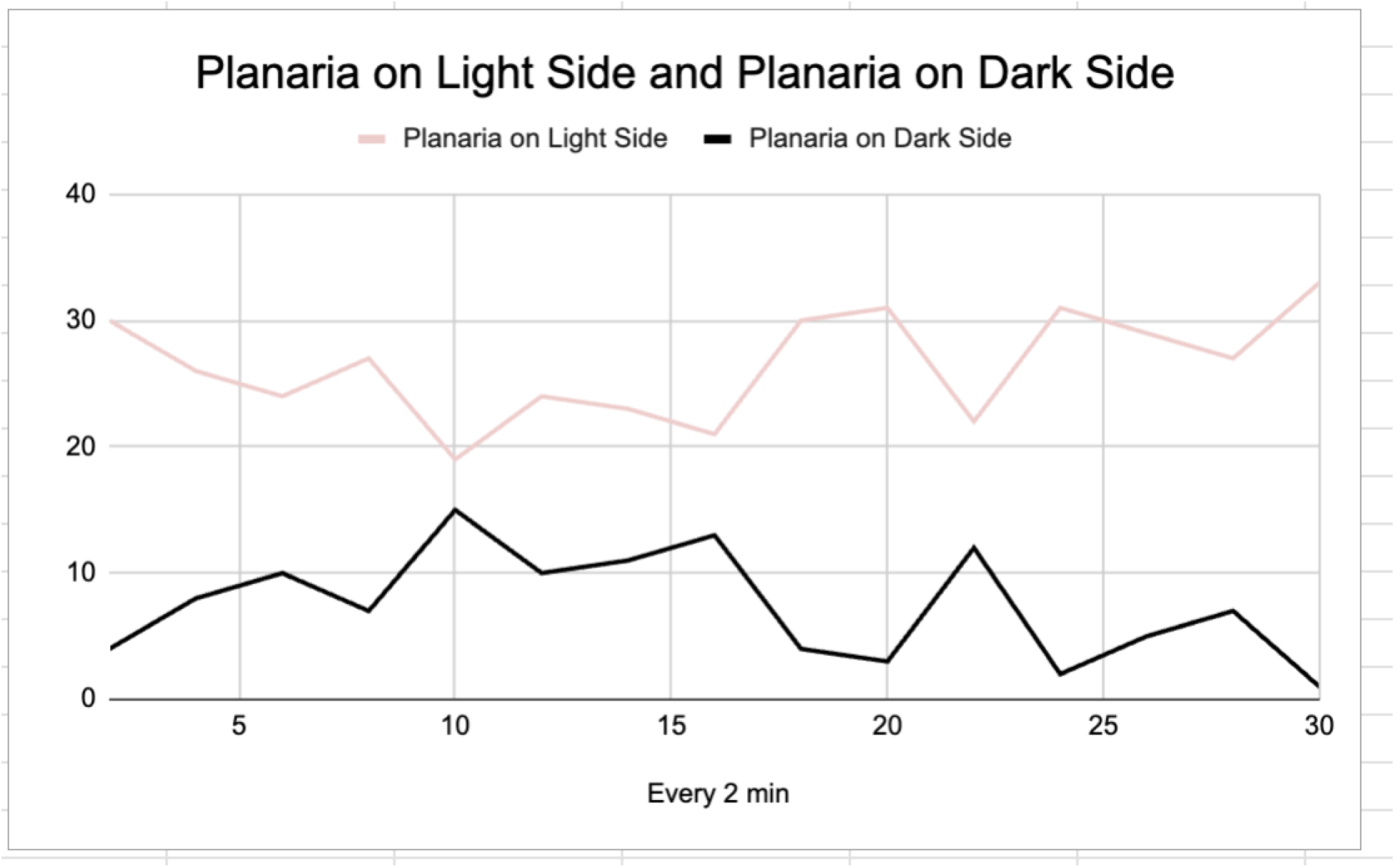
Post-training preference of planaria between the light and dark sides of the test arena, recorded every 2 minutes over 30 minutes (n = 34). After training, planaria showed a strong preference for the light side, demonstrating a learned behavioral shift compared to pre-training.

**Figure 3:**
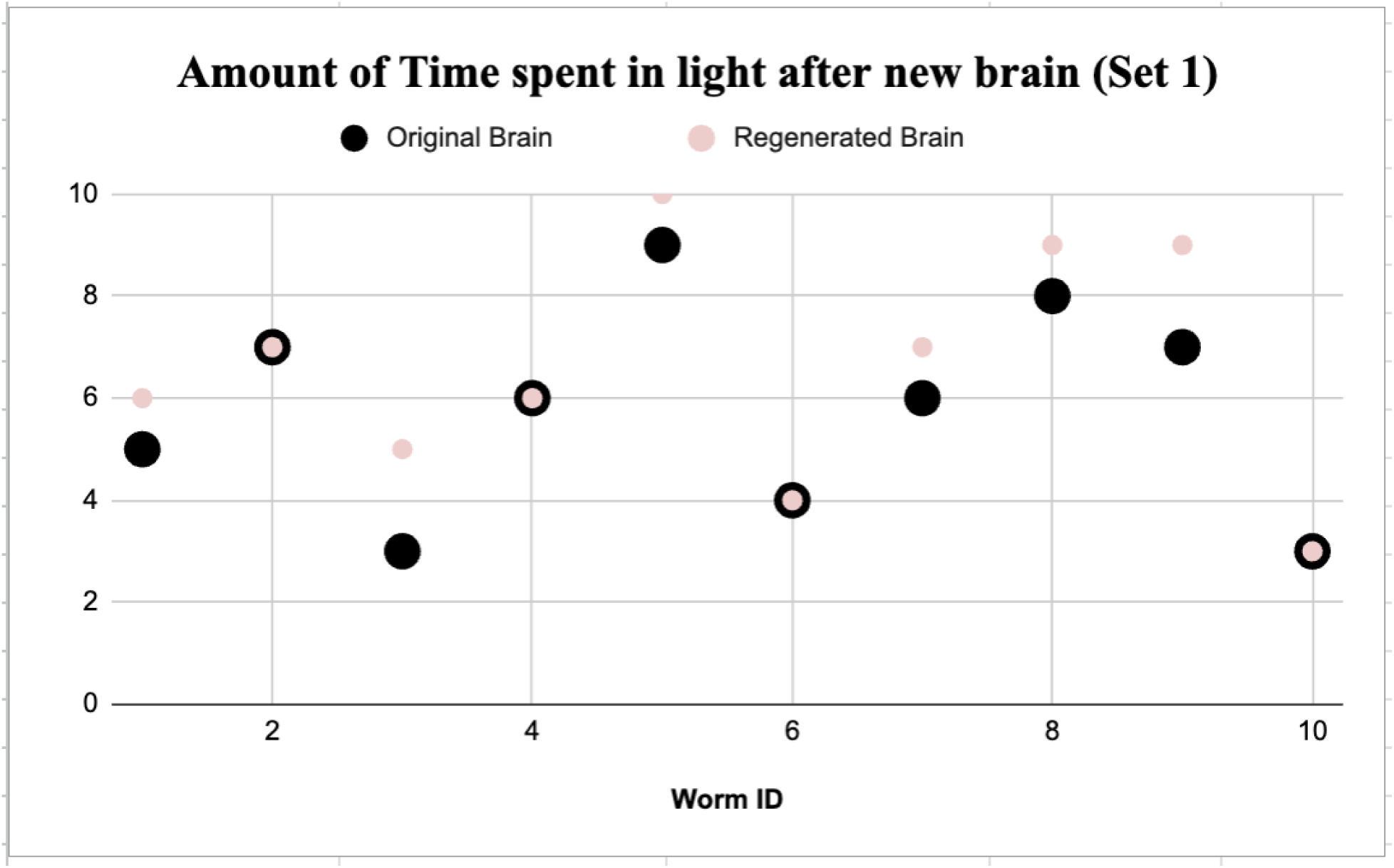
Amount of time spent in light by individual planaria before and after brain regeneration (Set 1). Black circles represent original brain trials; pink circles represent regenerated brain trials.

**Figure 4:**
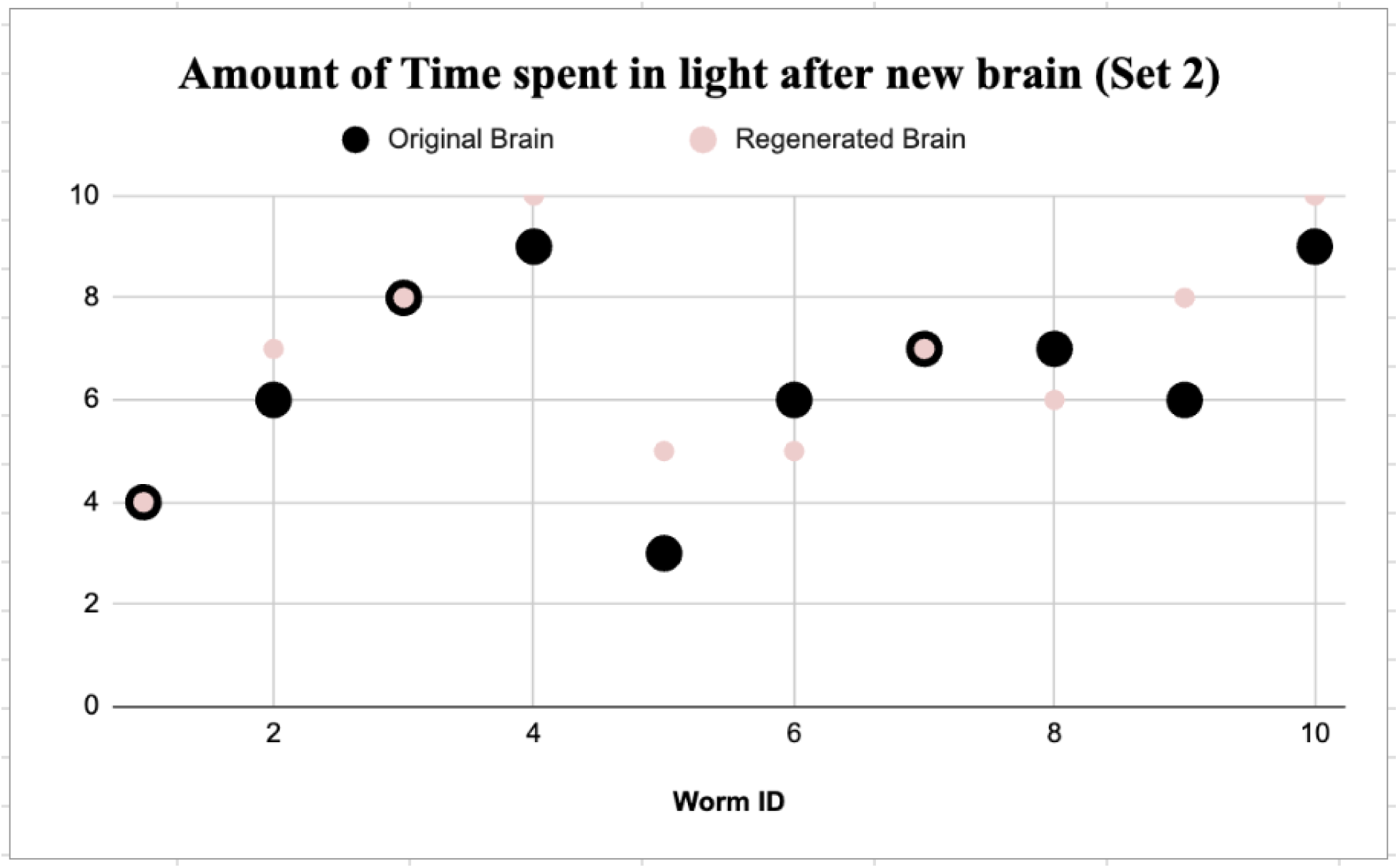
Amount of time spent in light by individual planaria before and after brain regeneration (Set 2). Black circles represent original brain trials; pink circles represent regenerated brain trials.

**Figure 5:**
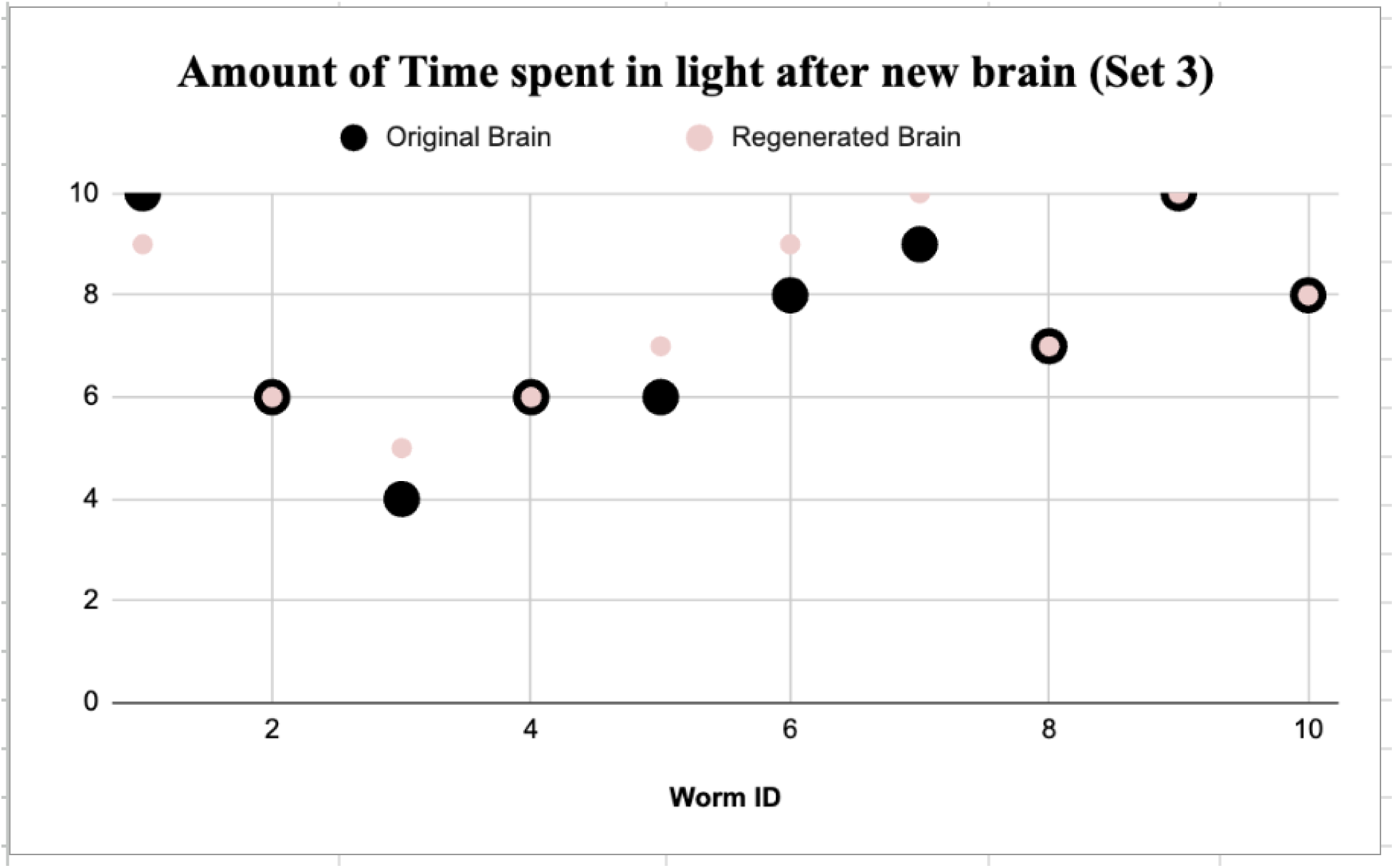
Amount of time spent in light by individual planaria before and after brain regeneration (Set 3). Black circles represent original brain trials; pink circles represent regenerated brain trials.

## Acknowledgments

We thank Wesley Clawson for helpful comments on the manuscript.

---

**Table A1.**
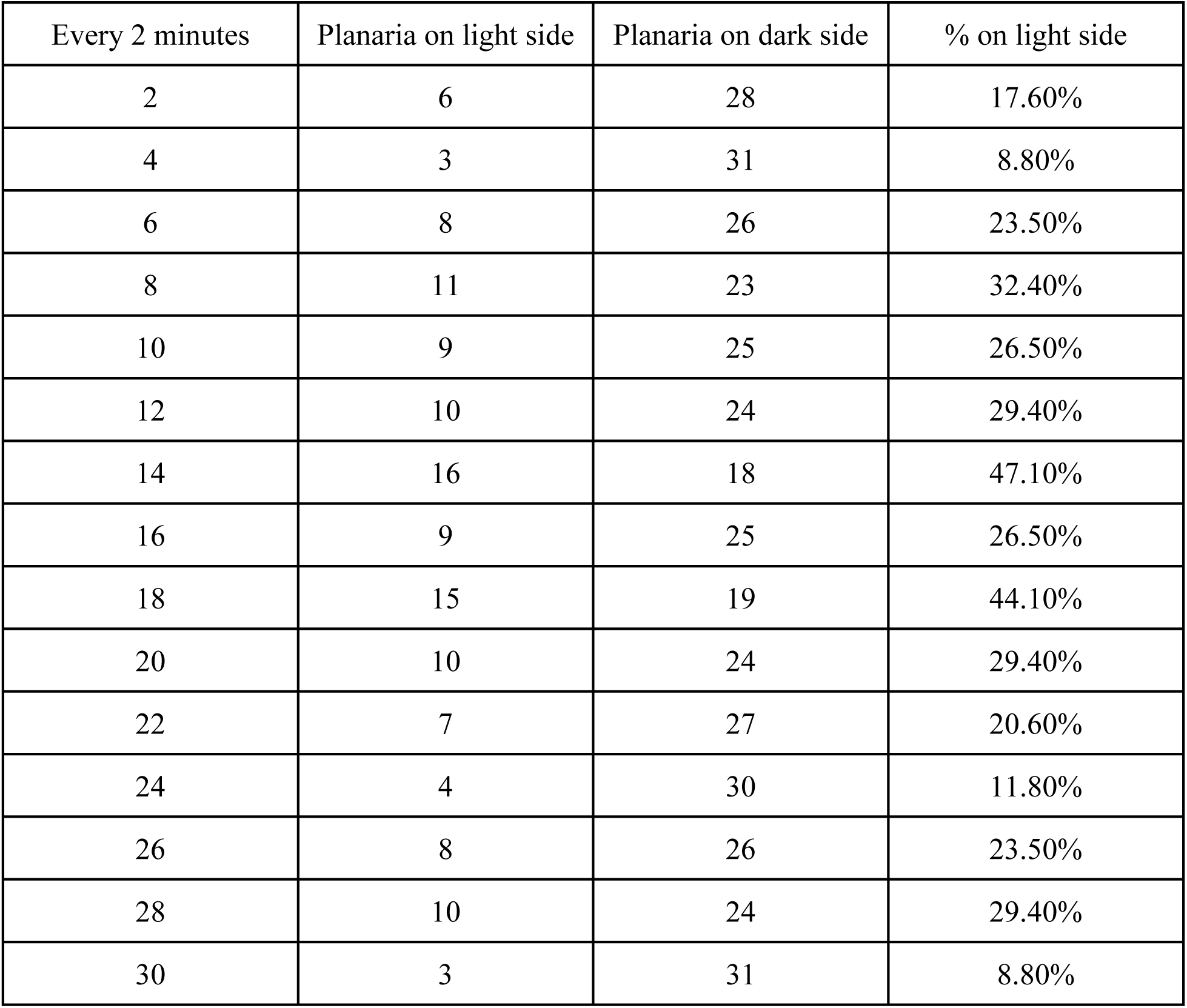
Planarian Light vs. Dark Side Preference (34 worms). Each entry shows the number of planaria observed on the Light side and Dark side at 2-minute intervals during a 30-minute free-movement trial. *Before Training* (baseline) indicates innate preference.

**Table A2.**
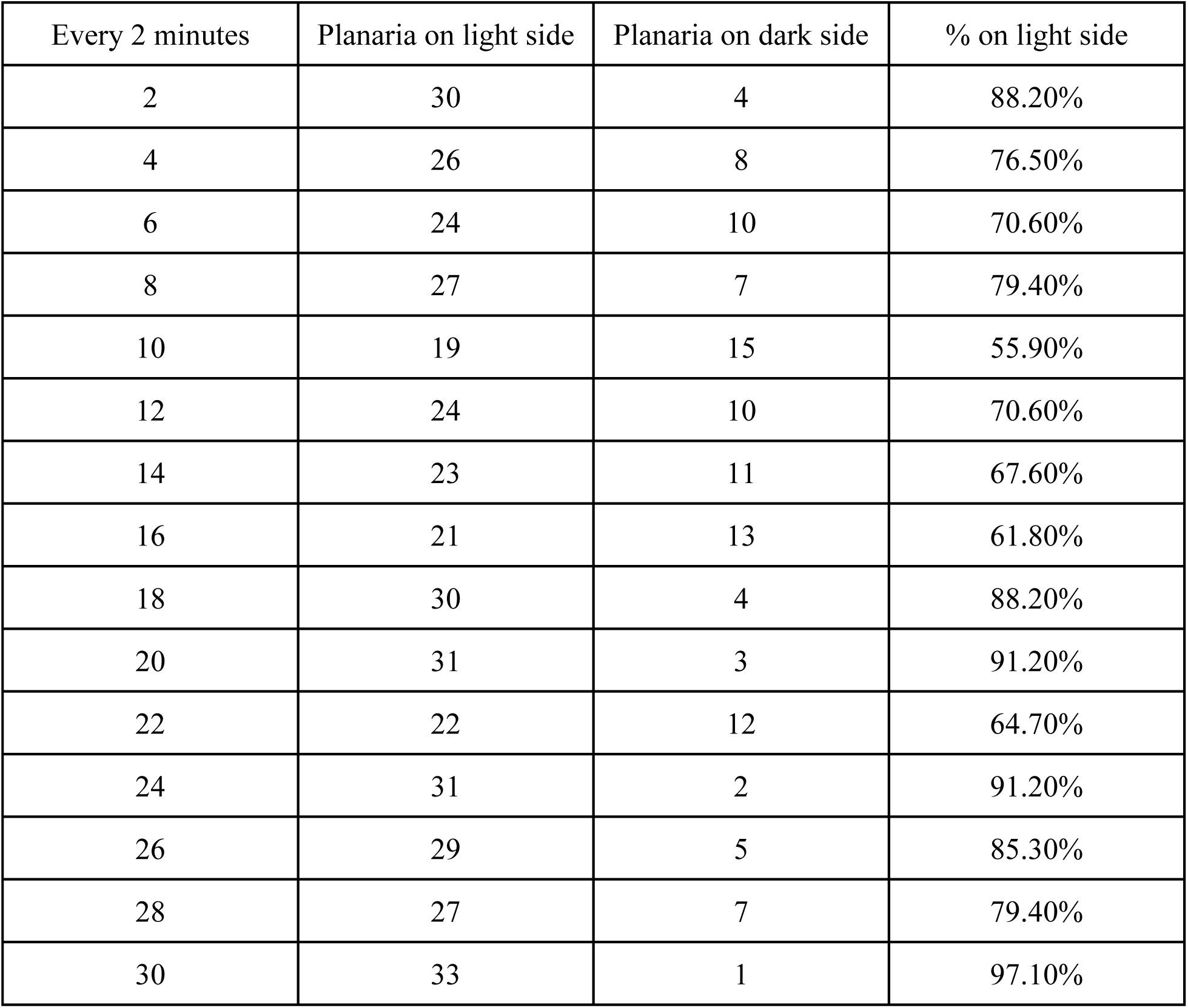
Planarian Light vs. Dark Side Preference After Training (34 worms). Each entry shows the number of planaria observed on the Light side and Dark side at 2-minute intervals during a 30-minute free-movement trial. *After training* shows the learned change in preference after conditioning with light-food pairing.

| Every 2 minutes | Planaria on light side | Planaria on dark side | % on light side |
| --- | --- | --- | --- |
| 2 | 30 | 4 | 88.20% |
| 4 | 26 | 8 | 76.50% |
| 6 | 24 | 10 | 70.60% |
| 8 | 27 | 7 | 79.40% |
| 10 | 19 | 15 | 55.90% |
| 12 | 24 | 10 | 70.60% |
| 14 | 23 | 11 | 67.60% |
| 16 | 21 | 13 | 61.80% |
| 18 | 30 | 4 | 88.20% |
| 20 | 31 | 3 | 91.20% |
| 22 | 22 | 12 | 64.70% |
| 24 | 31 | 2 | 91.20% |
| 26 | 29 | 5 | 85.30% |
| 28 | 27 | 7 | 79.40% |
| 30 | 33 | 1 | 97.10% |

## References

1. Saló, E., et al., Planarian regeneration: achievements and future directions after 20 years of research. Int J Dev Biol, 2009. 53(8-10): p. 1317–27.

2. Reddien, P.W., The Cellular and Molecular Basis for Planarian Regeneration. Cell, 2018. 175(2): p. 327–345.

3. Gentile, L., F. Cebria, and K. Bartscherer, The planarian flatworm: an in vivo model for stem cell biology and nervous system regeneration. Dis Model Mech, 2011. 4(1): p. 12–9.

4. Jacobson, A.L.a.M., James V., Research on learning in the planarian. Carolina Tips, 1962. XXV(7): p. 25–27.

5. McConnell, J.V., A.L. Jacobson, and D.P. Kimble, The effects of regeneration upon retention of a conditioned response in the planarian. Journal of Comparative Physiology and Psychology, 1959. 52: p. 1–5.

6. Deochand, N., M.S. Costello, and M.E. Deochand, Behavioral Research with Planaria. Perspectives on Behavior Science, 2018. 41(2): p. 447–464.

7. Blackiston, D., T. Shomrat, and M. Levin, The Stability of Memories During Brain Remodeling: a Perspective. Communicative & Integrative Biology, 2015. 8(5): p. e1073424.

8. Jacobson, A.L., C. Fried, and S.D. Horowitz, Planarians and memory. Nature, 1966. 209(23): p. 599–601.

9. Jacobson, A.L., Fried, C., and Horowitz, S.D., Planarians and memory: Transfer of learning by injection of ribonucleic acid. Nature, 1966. **209**: p. 599–601.

10. Wells, P.H., Training flatworms in a Van Oye maze, in Chemistry of Learning, W.C. Corning and S.C. Ratner, Editors. 1967, Plenum: New York. p. 251–254.

11. Verster, F.D.B. and J.T. Tapp, ON RNA AND MEMORY - TRANSFER OF LEARNED BEHAVIOR BY INJECTIONS OF RNA. Psychological Reports, 1967. 21(3): p. 937-C.

12. McConnell, J.V., The modern search for the engram, in A manual of psychological experimentation on planarians, J.V. McConnell, Editor. 1967, Journal of Biological Psychology: Ann Arbour.

13. Jacobson, A., S. Horowitz, and C. Fried, Classical conditioning, pseudoconditioning, or sensitization in the planarian. Journal of Comparative & Physiological Psychology, 1967. 64(1): p. 73–79.

14. Rilling, M., The mystery of the vanished citations: James McConnell’s forgotten 1Sc0s quest for planarian learning, a biochemical engram, and celebrity (vol 51, pg 58S, 1SSc). American Psychologist, 1996. 51(10): p. 1039–1039.

15. Corning, W.C. and D. Riccio, The planarian controversy, in Molecular approaches to learning and memory, W. Byrne, Editor. 1970, Academic Press: New York. p. 107–150.

16. Queenan, B.N., et al., On the research of time past: the hunt for the substrate of memory. Ann N Y Acad Sci, 2017. 1396(1): p. 108-125.

17. Gershman, S.J., et al., Reconsidering the evidence for learning in single cells. Elife, 2021. 10.

18. Langille, J.J. and C.R. Gallistel, Locating the engram: Should we look for plastic synapses or information-storing molecules? Neurobiol Learn Mem, 2020. **169**: p. 107164.

19. Gallistel, C.R., The physical basis of memory. Cognition, 2020: p. 104533.

20. Gallistel, C.R., Finding numbers in the brain. Philos Trans R Soc Lond B Biol Sci, 2017. 373(1740).

21. Gallistel, C.R., The Coding Question. Trends Cogn Sci, 2017. 21(7): p. 498–508.

22. Gallistel, C.R., Numbers and brains. Learn Behav, 2017. 45(4): p. 327–328.

23. Yassa, M.A. and C.E. Stark, Pattern separation in the hippocampus. Trends Neurosci, 2011. 34(10): p. 515–25.

24. Neuhof, M., M. Levin, and O. Rechavi, Vertically- and horizontally-transmitted memories - the fading boundaries between regeneration and inheritance in planaria. Biol Open, 2016. 5(9): p. 1177–88.

25. Shomrat, T. and M. Levin, An automated training paradigm reveals long-term memory in planarians and its persistence through head regeneration. The Journal of experimental biology, 2013. 216(Pt 20): p. 3799–810.

26. Blackiston, D.J., E. Silva Casey, and M.R. Weiss, Retention of memory through metamorphosis: can a moth remember what it learned as a caterpillar? PLoS One, 2008. 3(3): p. e1736.

27. Alloway, T.M., Retention of Learning through Metamorphosis in Grain Beetle (Tenebrio-Molitor). American Zoologist, 1972. 12(3): p. 471–472.

28. Ray, S., Survival of olfactory memory through metamorphosis in the fly Musca domestica. Neuroscience Letters, 1999. 259(1): p. 37–40.

29. Sheiman, I.M., E.F. Balobanova, and N.D. Kreshchenko, Regulation of development of the grain beetle Tenebrio molitor by neuropeptides. Invertebrate Reproduction & Development, 1999. 36(1-3): p. 105–110.

30. Sheiman, I.M. and K.L. Tiras, Memory and morphogenesis in planaria and beetle, in Russian contributions to invertebrate behavior, C.I. Abramson, Z.P. Shuranova, and Y.M. Burmistrov, Editors. 1996, Praeger: Westport, CT. p. 43–76.

31. Bedecarrats, A., et al., RNA from Trained Aplysia Can Induce an Epigenetic Engram for Long-Term Sensitization in Untrained Aplysia. eNeuro, 2018. 5(3).

32. Chen, L., et al., Adult hippocampal neurogenesis: New avenues for treatment of brain disorders. Stem Cell Reports, 2025. 20(9): p. 102600.

33. Li, X., X. Bao, and R. Wang, Neurogenesis-based epigenetic therapeutics for Alzheimer’s disease (Review). Mol Med Rep, 2016. 14(2): p. 1043–53.

34. Uwishema, O., et al., Stem cell therapy use in patients with dementia: a systematic review. Int J Emerg Med, 2025. 18(1): p. 95.

35. Fields, C., J. Bischof, and M. Levin, Morphological Coordination: A Common Ancestral Function Unifying Neural and Non-Neural Signaling. Physiology, 2020. 35(1): p. 16–30.

36. O’Brien, T., et al., Machine learning for hypothesis generation in biology and medicine: exploring the latent space of neuroscience and developmental bioelectricity. Digital Discovery, 2024. 3(2): p. 249–263.

37. Chen, S., et al., Reinstatement of long-term memory following erasure of its behavioral and synaptic expression in Aplysia. eLife, 2014. 3: p. e03896.

38. Glanzman, D.L., New tricks for an old slug: the critical role of postsynaptic mechanisms in learning and memory in Aplysia. Prog Brain Res, 2008. **169**: p. 277–92.

39. Nahm, M., et al., Terminal lucidity: a review and a case collection. Arch Gerontol Geriatr, 2012. 55(1): p. 138–42.

40. Mukherjee, D., et al., Salient experiences are represented by unique transcriptional signatures in the mouse brain. Elife, 2018. 7.

41. Dent, E.W., Of microtubules and memory: implications for microtubule dynamics in dendrites and spines. Mol Biol Cell, 2017. 28(1): p. 1–8.

42. Craddock, T.J., J.A. Tuszynski, and S. Hameroff, Cytoskeletal signaling: is memory encoded in microtubule lattices by CaMKII phosphorylation? PLoS computational biology, 2012. 8(3): p. e1002421.

43. Levin, M., Self-Improvising Memory: A Perspective on Memories as Agential, Dynamically Reinterpreting Cognitive Glue. Entropy (Basel), 2024. 26(6).

44. Wagner, D.E., I.E. Wang, and P.W. Reddien, Clonogenic neoblasts are pluripotent adult stem cells that underlie planarian regeneration. Science, 2011. 332(6031): p. 811–6.

